# Chemical inhibition of Exon Junction Complex assembly impairs mRNA localization and neural stem cells ciliogenesis

**DOI:** 10.1101/2025.02.25.640098

**Authors:** Tommaso Villa, Oriane Pourcelot, David Dierks, Marion Faucourt, Cindy Burel, Floric Slimani, Léa Guyonnet, Nathalie Spassky, Schraga Schwartz, Edouard Bertrand, Olivier Bensaude, Hervé Le Hir

## Abstract

The Exon Junction Complex (EJC) is formed by the essential eIF4A3, MAGOH and Y14 core proteins. It is universally deposited during splicing at exon-exon junctions. The EJC is known to impact almost every post-transcriptional regulatory step throughout the life of mRNAs including their modifications, splicing, decay and trafficking. Its dysregulation leads to neurodevelopmental pathologies. Here we show that EJC-i, a compound known to block the ATPase activity of eIF4A3, inhibits *de novo* EJC assembly. EJC-i and targeted knockdown of either eIF4A3 or Y14 core EJC subunits lead to very similar phenotypes by impacting the destiny of mRNAs due to alterations in alternative splicing, nonsense-mediated mRNA decay, genome-wide m6A methylation and proper intracellular addressing of specific transcripts, in particular to the centrosome. Both EJC impairment methods disrupt the centrosome function that might be responsible for mitotic arrest at prometaphase. As a small molecule that readily diffuses into cells, EJC-i is a particularly attractive easy to use and versatile tool to investigate EJC functions in live cells or whole organism that are not prone to genetic manipulation. Indeed, this property was used to disrupt ciliogenesis in primary neural stem cells.

## INTRODUCTION

Regulation of gene expression operates at multiple and interconnected levels to ensure the proper flow of genetic information. The modulation of regulatory networks in response to environmental cues and internal stimuli is essential for the homeostasis of living organisms. Indeed, dysregulation of gene expression mechanisms often results in diseases. These mechanisms act at many levels all along the path leading from the information stored in the genome to the production of proteins, therefore encompassing epigenetic, transcriptional, post-transcriptional, translational and post-translational processes. Loss of regulation of post-transcriptional control has gained attention as a pathogenic mechanism because of the identification of a large number of pathogenic mutations in ubiquitously expressed RNA binding proteins (RBPs) (1). These RBPs cover mRNAs from the moment they emerge from RNA polymerase II and all along their life within a cell, and partake into many post-transcriptional regulatory steps leading to functional and mature molecules (2). Further, the repertoire of RBPs on individual transcripts evolves with each step of a mRNA life, and this dynamic composition of the mRNA ribonucleoparticles (mRNPs) determines their specific pattern of processing, intra-cellular localization, translation and decay (2, 3).

Association of proteins with mRNAs is either dictated by defined cis-acting features of the RNA molecules, such as specific sequences or structures, or result from the history of mRNA processing. The Exon Junction Complex (EJC) is a paradigmatic example of this second class of RBPs (4, 5). The EJC is a multiprotein complex marking mRNAs that have not yet been translated. It is deposited during the splicing reaction, 27 nucleotides upstream of exon-exon junctions (6, 7). The core of the complex is composed by the three proteins eIF4A3/DDX48, MAGOH and Y14/RBM8A (8). It interacts with accessory proteins such as MLN51/CASC3/BARENTSZ or the ASAP and PSAP complexes that comprise RNPS1, SAP18 and ACIN1 or PNN, respectively. The DEAD-box RNA helicase eIF4A3 clamps the RNA by binding to the sugar-phosphate backbone of mRNA independently of the sequence, and is stabilized and locked onto the RNA by the heterodimer MAGOH/Y14 (9, 10). Recent evidence suggests a universal deposition of EJC (7). However, a possible specialization of the EJC may occur depending on transcripts and/or exon junctions and accessory protein composition (11–13). EJCs are disassembled by translation-dependent or translation-independent mechanisms and their persistence onto mRNA is variable and transcript-specific (14, 15).

Once clamped on mRNA, the EJC core complex acts as a binding platform for several peripheral factors first in the nucleus and, after mRNA export, in the cytoplasm. Together, these factors are involved in post-transcriptional events including splicing, nuclear export, translation initiation, Nonsense Mediated Decay (NMD) (12, 16–20). A function of the EJC in the localization of some mRNAs to specific subcellular departments has been solidly established, although limited to two cases. The first example described was the *oskar* mRNA that is transported from nurse cells to the posterior pole in *D. melanogaster* oocytes (21). More recently, we identified the *NIN* mRNA, coding for the Ninein core component of centrosomes as a transcript whose localization to centrosomes relies on the EJC in quiescent RPE1 cells (22). Therefore, EJC-dependent RNA localization is present in vertebrates as well. Furthermore, the EJC contributes in excluding m6A RNA modification around spliced junctions (23–26).

The EJC central role in packaging the evolving mRNPs is reflected by the fact that an altered dosage of its components leads to pathological states such as cancer or developmental defects (27). In particular, the EJC is essential for neural stem cells division, differentiation and brain development (27, 28).

In order to understand the mechanisms by which the EJC participates in the wide array of regulatory steps it has been linked to, the ability to modulate its activity becomes a key issue. However, akin to other key cellular regulatory factors, all three core subunits of the EJC are encoded by essential genes (29), therefore precluding their knockout in cells. So far, blocking EJC activity relies on the core EJC proteins knockdown (KD), either using siRNAs (16, 17, 20, 22, 30–34), or using dTAG-mediated degradation of the EJC core subunit Y14 in CRISPR/Cas9 edited cell lines (24). However, KD approaches remain limited as they are slow processes, from several hours for dTAG up to 2-3 days for siRNAs and difficult to control. As a consequence, indirect effects may occur.

Chemical inhibitors are particularly attractive because they diffuse rapidly into cells (minutes) and can be used in different cell types and potentially, in whole organisms. A 1, 4-diacylpiperazine derivative (which we will refer to from now on as EJC-i for EJC-inhibitor) has recently been identified as a selective inhibitor of eIF4A3 ATPase activity (35). It has been shown to alter gene expression, NMD and the cell cycle (33, 36, 37). However, its mechanism of action remains poorly understood. In this study, we show that EJC-i blocks *de novo* EJC assembly both *in vitro* and *in vivo*, impairing a wide spectrum of established EJC-dependent functions. We exploit the extreme flexibility and ease of application of the inhibitor to identify novel EJC-dependently localized mRNAs and to strengthen EJC implication in centrosome function and ciliogenesis in human cells.

## MATERIAL AND METHODS

### Cell culture and treatments

All cells were maintained in DMEM high glucose (Dutscher) supplemented with 10% of fetal bovine serum (Sigma-Aldrich), and 1% of penicillin and streptomycin (Sigma-Aldrich) at 37°C. Cell lines used in this study were derivative of HEK293T or HeLa. Construction of HEK293T Y14-HA-dTAG was described (24), and the HEK293T eIF4A3-HA-dTAG was obtained following the same procedure. Construction of cell line OB9, co-expressing MAGOH-LgBiT and FLAG-eIF4A3-SmBiT, has been described (14). HeLa cells expressing Centrin1-GFP have been previously described (48). Cells were cultivated at 70% confluency and treated with DMSO 1:500, or the indicated dose (see Results) ranging from 1 to 20 µM EJC-i (MedChemExpress) in DMSO, or 50 nM dTAG13 or dTAG1 (Sigma-Aldrich) in DMSO, and harvested after the indicated time post-treatment.

### Luciferase assays

These assays were performed essentially as described (14). 100 ng of furimazine, 1 mg/ml stock solution in 50:50 ethanol/propylene glycol) in 30 μl of 0.5x Passive Lysis Buffer (Promega #E1941) per assay were added to 50 μl of lysates or cell suspensions. The luciferase activity was detected with a Berthold TriStar LB941 luminometer. For kinetic experiments, OB9 cells were seeded on polylysine-coated 12-well plates and EJC-i was added at defined times prior to lysis. Incubation was arrested on ice, the medium was rapidly sucked off, and cells were scraped in 400 μl ice-chilled 0.5x Passive Lysis Buffer per well. Each measure is an average for three wells (technical replicates) treated independently. Three wells per plate were exposed to DMSO 1:500 and their average luciferase activity used to normalize the data for drug-treated cells on the same plate. Standard deviations are determined from biological replicates obtained on different days.

### Western blot and co-immunoprecipitation

For coimmunoprecipitation, 10-cm plates of confluent cells were collected and washed in 1 ml PBS. Cell were lysed for 15 min at 4 °C in 300 μl of lysis buffer (20 mM Tris-HCl, pH 7.5, 150 mM NaCl, 1 mM Na2EDTA, 1 mM EGTA, 1% NP-40, 1% sodium deoxycholate, 1% protease-inhibitor mixture), with 10 U RNase T1 (Fermentas) and 12 U RQ1 DNase (Promega). Cellular debris was removed by centrifugation at 13000 rpm for 10 min. The lysates were incubated with 20 μl protein A-coupled Dynabeads (Life Technologies) with or without antibody for 2 h at 4 °C. The beads were washed extensively with IP buffer 150 (10 mM Tris-HCl, pH 7.5, 150 mM NaCl, 2.5 mM MgCl_2_, 1% NP-40, 1% protease-inhibitor mixture), and the bound proteins were eluted with Laemmli buffer. Total proteins, input proteins or eluted proteins were resolved by 4-12% SDS-PAGE and electrotransferred to nitrocellulose membrane (Schleicher & Schuell). Membranes were blocked in PBS containing 5% nonfat dry milk and 0.05% Tween 20. Rabbit polyclonal anti-GAPDH, anti-Y14, anti-MAGOH and anti-eIF4A3 were used as primary antibodies at 1:1000 dilution. After washing, membranes were incubated with secondary antibodies: either goat anti-rabbit-HRP or Clean-Blot™ IP Detection Reagent (HRP) (Thermofisher). Protein-antibody complexes were visualized by an enhanced chemiluminescence detection system (SuperSignal West Pico Plus, Thermo Scientific).

### EJC reconstitution assay

These assays were performed essentially as described (38). Proteins (2 µg of each) were mixed in binding buffer (BB-125) containing 20 mM HEPES (pH 7.5), 125 mM NaCl, 1 mM magnesium diacetate, 1 mM DTT, 5% (v/v) glycerol and 0.1% (w/v) NP-40, complemented or not with 2 mM ADPNP, 1-20 µM EJC-i or 1.6% DMSO, and 0.5 µM biotinylated ssRNA in a final volume of 60 µl. After 20 min at 30°C, 5 µl of magnetic streptavidin beads (MyOne, Dynal) and 140 µl of BB-250 (250 mM NaCl) were added. After rotation for 1 h at 4°C, beads were washed three times with 500 µl BB-250 and then eluted with Laemmli buffer, boiled and loaded on 4-12% SDS-PAGE along with inputs and with a protein marker (broad range, New England Biolabs). Proteins were visualized by Coomassie staining.

### RNA extraction, Reverse Transcription, End-Point and Quantitative RT-PCR

Total RNA was extracted from cells using TRI reagent (Ambion) according to manufacturer’s protocol. The RNA was digested with 5 U RQ1 DNase (Promega) for 1 h at 37°C before phenol extraction and precipitation. Reverse transcription was performed using 1 µg RNA with random hexamers and Superscript IV reverse transcriptase (Invitrogen) according to manufacturer’s protocol. After heat inactivation, samples were treated with 5 U RNase H for 30 min at 37°C. For end-point PCRs, 1% cDNA was used as template using GoTaq G2 Flexi (Promega) and 0.2 µM final concentration of sense and antisense primer. After 30 PCR cycles, the samples were resolved by electrophoresis on ethidium bromide-stained, 2% agarose TBE gels and visualized by trans-UV illumination. For qPCRs, 1% cDNA was used as template using PowerUP SYBR Green Master Mix (Applied Biosystems) on CFX384 Real time system (Bio-Rad). Quantification was performed using the ΔΔCt method with RNA levels normalized by housekeeping gene GAPDH. Controls without reverse transcriptase were systematically run in parallel to estimate the contribution of contaminating DNA. Amplification efficiencies were calculated for every primer pair in each amplification reaction.

### m6A-seq Library Preparation

Whole cell RNA from the samples in the EJC-i time course experiment was poly-A selected using oligo dT-beads (Dynabeads mRNA DIRECT Kit). A total of 200 ng of poly-A selected mRNA per sample was used to prepare m6A-seq2 libraries following the step-by-step protocol, as previously described (47). The m6A-IP and Input NGS libraries were sequenced paired-end on the NovaSeq 6000 platform.

### m6A-seq2 Data Processing and Analysis

Sequencing reads were aligned to the hg19 human reference genome using STAR/2.7.9a. The bam alignment files were filtered to include only unique alignments, and gene coverage was extracted using the bam2Endreads R script (47) based on the canonical hg19 UCSC gene annotation. m6A gene-level (m6A-GI) estimation was performed as described (47). Meta-analysis of m6A coverage for long internal exons (excluding the first or last exon and those >600 nucleotides) and last exons (for genes with >1 exon and last exons >400 nucleotides) was conducted. For the analysis, m6A enrichment scores (m6A-IP/Input coverage) were calculated for each gene with sufficient coverage in 20 nt windows. The median m6A enrichment score for each sample was analyzed, and the mean of the replicates was normalized by the mean of the control timepoint (0 h) to calculate the meta m6A enrichment score.

### De Novo m6A Peak Calling

De novo m6A peak calling was performed for each sample as described (70). From all detected peaks, high-confidence m6A sites were defined as de novo m6A peaks in which the summit was within five bases of the nearest adenosine in a DRACH motif context. m6A site score calculation (m6A-IP/Input coverage) was performed for the detected high-confidence m6A sites across all samples.

### Quantification and Statistical Analysis

All statistical analyses were performed using R 4.1.0. Details of the analyses are provided in the figure legends and result sections. Figures were prepared using basic R packages and ggplot2.

### Flow cytometry analysis

Cells were trypsinized and washed in D-PBS. Following the last centrifugation, cells were fixed and permeabilized by resuspension in cold 70% EtOH. They were incubated for 48h at -20°C. Cells were washed in DPBS and counted. Equal number of cells per condition were labeled by 10 μg/ml of HOECHST 33342 in the presence of 100 μg/ml RNase A at 4°C for 48 hours. After HOECHST 33342 staining, samples were analyzed on a ZE5 cell analyzer (Bio-Rad) using Everest acquisition software (Version 2.5). Data were analyzed using FlowJo software (FlowJo, LLC, version 10.8.1). Briefly, cells were selected based on their morphology (SSC-A versus FSC-A: “Cells” population) then doublets of cells were excluded based on DNA content parameters (Hoechst-A versus Hoechst-W: “Single cells” population). Cell cycle analysis was performed on “Single cells” population using FlowJo module for cell cycle.

### smFISH

Probe sets of DNA oligonucleotides targeting the mRNAs of interest were designed based on the Oligostan script(71), and used as previously described (71). RNA probes were obtained as described previously (48). For RNA probes, 50 ng of unlabelled RNA probes and 25 ng of each fluorescent secondary probes (TYE-650-labeled LNA oligonucleotides, Qiagen) were prehybridized in 100µl of a solution containing 7.5 M urea (Sigma-Aldrich), 0.34 mg/ml tRNA (Sigma-Aldrich) and 10% Dextran sulfate. The pre-hybridization was performed in a thermocycler, with the following program: 90 °C for 3 min, 53 °C for 15 min, forming fluorescent duplexes. Cells were grown in a 96-well plate with glass bottom (PhenoPlate, Revvity), fixed with 4% PFA at RT and permeabilized with 70% ethanol overnight at 4 °C. They were washed with PBS and incubated 15 min at RT in the hybridization buffer (1 x SSC, 7.5 M urea) before the addition of the 100µl of fluorescent duplexes in each well. Hybridization was performed at 48 °C overnight. The next day, the plate was washed 8 times for 20 min each in 1 x SSC, and 7.5 M urea at 48 °C. Finally, cells were washed 3 times with PBS at RT for 10 min each, stained with 5 µg/mL DAPI, and mounted in 90% glycerol (VWR), 1 mg/mL p-Phenylenediamine (Sigma-Aldrich), PBS pH 8.

### Imaging

smFISH imaging for HeLa Centrin-GFP cells was performed on an Opera Phenic High-Content Screening System (Revvity), with a 63x water-immersion objective (NA 1.15). The imaging on SunTag cell lines was performed on a Leica thunder with a 63x oil-immersion objective (NA 1.4). Three-dimensional images were acquired, with a z-spacing of 0.6 µm.

### Image analysis

Quantifications were based on an image analysis pipeline including cell segmentation, threshold-based spot detection, dense region deconvolution and density-based clustering algorithm (DBSCAN). Firstly, nuclei and cytoplasm segmentation are performed in 2D on image stack mean projection using neural network Cellpose (72). Secondly, single molecules are detected as 3D spots using the Big-FISH package (48) (https://github.com/fish-quant/big-fish) which relies on an intensity Laplacian of Gaussian (LoG) filter followed with a local maxima filter and a user adjusted thresholding. During analysis we were careful that same thresholds were used on stacks targeting the same mRNA. As local maxima detection can lead to a miss estimation of single molecule number in bright region a dense region deconvolution step is added (48). It consists in fitting the median detected spot with a 3D-gaussian to construct a reference spot. Then an estimation of single molecule number is done by reconstructing the bright region signal using the reference spot. Thirdly, a DBSCAN clustering algorithm is used to quantify foci. Finally, Big-FISH spot detection was used with the Centrin-GFP channel to detect centrosomes. The brightest candidate spot was considered as the centrosome, and we allowed cells to have a second centrosome if its intensity was lying in a ten percent range of the first centrosome. Normalized distances from the detected RNA spot to centrosome were then computed. In each cell, distances from the RNA spots to the closest centrosome were normalized using distances from a random RNA distribution. A distance inferior to 1 indicates centrosomal enrichment whereas a distance superior to one indicates anti-colocalization. The same calculation was done for the normalized distance to the cell membrane.

## RESULTS

### EJC-i decreases the amounts of EJC in live cells

Previous studies showed that EJC-i inhibits eIF4A3 ATPase and helicase activities *in vitro* (33, 35). However, EJC-i was not found to affect eIF4A3 ability to interact with EJC core components and to form an EJC (33). Yet, EJC-i impairs eIF4A3-mediated NMD *in vivo* (33, 36). To test the impact of EJC-i on EJC assembly in live cells, we coimmunoprecipitated EJC components from HEK293T cells treated with 20 µM for 4 hours, a non-toxic dose of EJC-i. Western blot analysis showed that EJC-i treatment reproducibly resulted in a reduction of the amount of both Y14 (52.2 ± 20.4 %) and MAGOH (65.5 ± 10.5 %) EJC subunits coimmunoprecipitated with eIF4A3 (Figure 1a, compare lanes 8 and 9). Because inhibition of EJC could derive from destabilization of eIF4A3, we monitored the steady-state levels of this core component of the complex after incubation for different times up to 24 hours in wild type HEK293T cells. 20 µM EJC-i did not alter eIF4A3 concentration even after long exposure (Supplementary Figure 1a).

**Figure 1.**
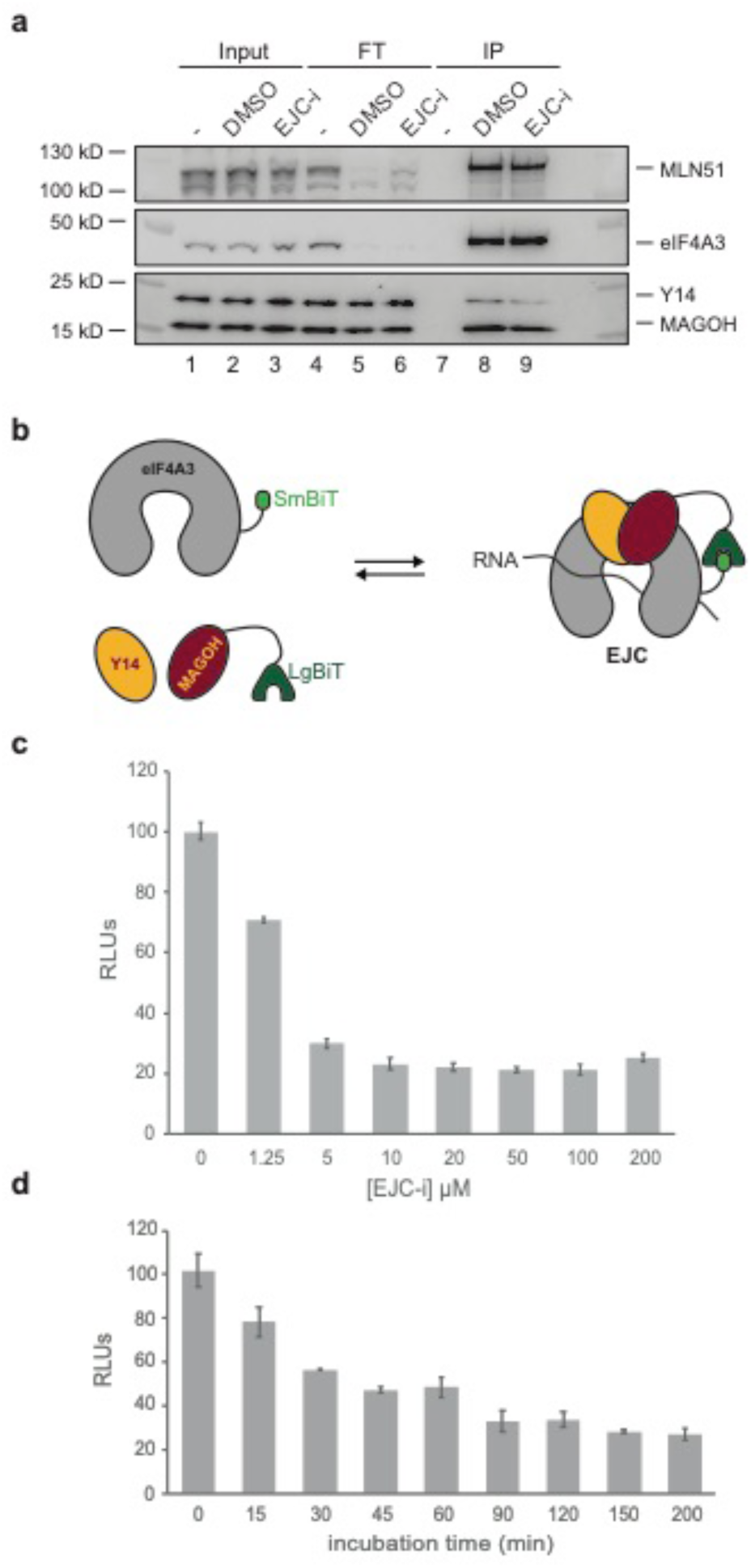
EJC-i decreases the amounts of EJC in live cells. **a** Western blot analysis of EJC proteins coimmunoprecipitated (IP) with anti-eIF4A3 from protein extracts of HEK293T cells treated with either DMSO or a 20 µM dose of EJC-i for 4 hours. Lanes labelled - represent no antibody controls. FT is the flow-through fraction. **b** Scheme depicting the EJC-NanoBiT system. **c** Response of a stable HEK293T cell line co-expressing MAGOH-LgBiT and FLAG-eIF4A3-SmBiT to a 2 hours incubation with the indicated µM doses of EJC-i expressed as relative luciferase units (RLUs). **d** Response kinetics measured as relative luciferase units (RLUs) of the same cell line as in (**c**) incubated with a 20 µM dose of EJC-i for the indicated times (minutes).

As Western blots are poorly quantitative, we turned to a highly quantitative split-luciferase assay that we have recently established to monitor assembly and disassembly rates of the EJC in live cells (14). This assay uses a stable cell line co-expressing MAGOH-LgBiT and FLAG-eIF4A3-SmBiT proteins where N-terminal (LgBiT) and C-terminal (SmBiT) moieties of nanoluciferase are fused to MAGOH and eIF4A3, respectively. Assembly of EJCs reconstitutes nanoluciferase (NanoBiT) activity (14) (Figure 1b). The split-luciferase cell line was incubated for 2 hours with different doses of EJC-i or DMSO as control. EJC-i reduced significantly the observed luciferase activity, reflecting a decrease in assembled EJC that reached an 80% plateau between 10 and 20 µM concentrations (Figure 1c). The effect of EJC-i was rapid and reached 50% inhibition within the first hour of incubation for 20 µM concentration (Figure 1d). Since the split-luciferase assays suggested a significant reduction in the amount of assembled recombinant EJC following EJC-i treatment, a Western blot analysis was performed on the cell line co-expressing MAGOH-LgBiT and FLAG-eIF4A3-SmBiT. Following immunoprecipitation with an anti-FLAG antibody and Western blot analysis of EJC components, reduction in the coimmunoprecipitated fraction of Y14 and both endogenous and LgBiT-tagged MAGOH proteins was observed (Supplementary Figure 1b, compare lanes 8 and 9).

Altogether, these results show that an EJC-i treatment decreases the amounts of assembled EJC in live cells.

### EJC-i inhibits *de novo* EJC assembly

In order to test more directly the effect of EJC-i on the EJC, we used an established *in vitro* EJC assembly assay (38). Briefly, recombinant EJC proteins (eIF4A3, MAGOH/Y14 and the Selor domain of MLN51) were first mixed with a 3’-end biotinylated ssRNA and the non-hydrolyzable ATP analog ADPNP before RNA precipitation with streptavidin beads and analysis of RNA-bound proteins on SDS-PAGE (38). *In vitro* EJC reconstitutions were performed in the presence or absence of EJC-i, or DMSO as control (Figure 2a). The complex was formed and the proteins efficiently coprecipitated only when ADPNP was included in the incubation (Figure 2b, compare lanes 7 and 8). Remarkably, incubation in the presence of 20 µM EJC-i resulted in complete loss of protein coprecipitation, indicative of inhibition of complex assembly (Figure 2b, compare lanes 9 and 10). *In vitro*, a stable binding of the EJC core to RNA requires its four protein components (38). Although much less stable, the interaction of eIF4A3 with RNA can be detected alone or in the presence of one of its partners. Thus, we also tested the effect of EJC-i on eIF4A3 individual interactions with the EJC core components separately. Whether with RNA, MLN51-Selor or the MAGOH/Y14 heterodimer, incubation in the presence of EJC-i always impaired eIF4A3 binding to RNA (Supplementary Figure 2a, 2b, 2c). Finally, we tested decreasing concentrations of EJC-i. Protein coprecipitation progressively increased with lowering the amount of inhibitor from 20 µM to 1 µM, indicating that inhibition of EJC assembly was dose-dependent, reaching its maximum at 20 µM (Figure 2c). To assess whether EJC-i impacts the stability of already formed EJC, EJC-i or DMSO were added to preassembled EJC coprecipitated on streptavidin beads. Notably, in this case, no effect of EJC-i was observed (Figure 2b, compare lanes 11 and 12).

**Figure 2.**
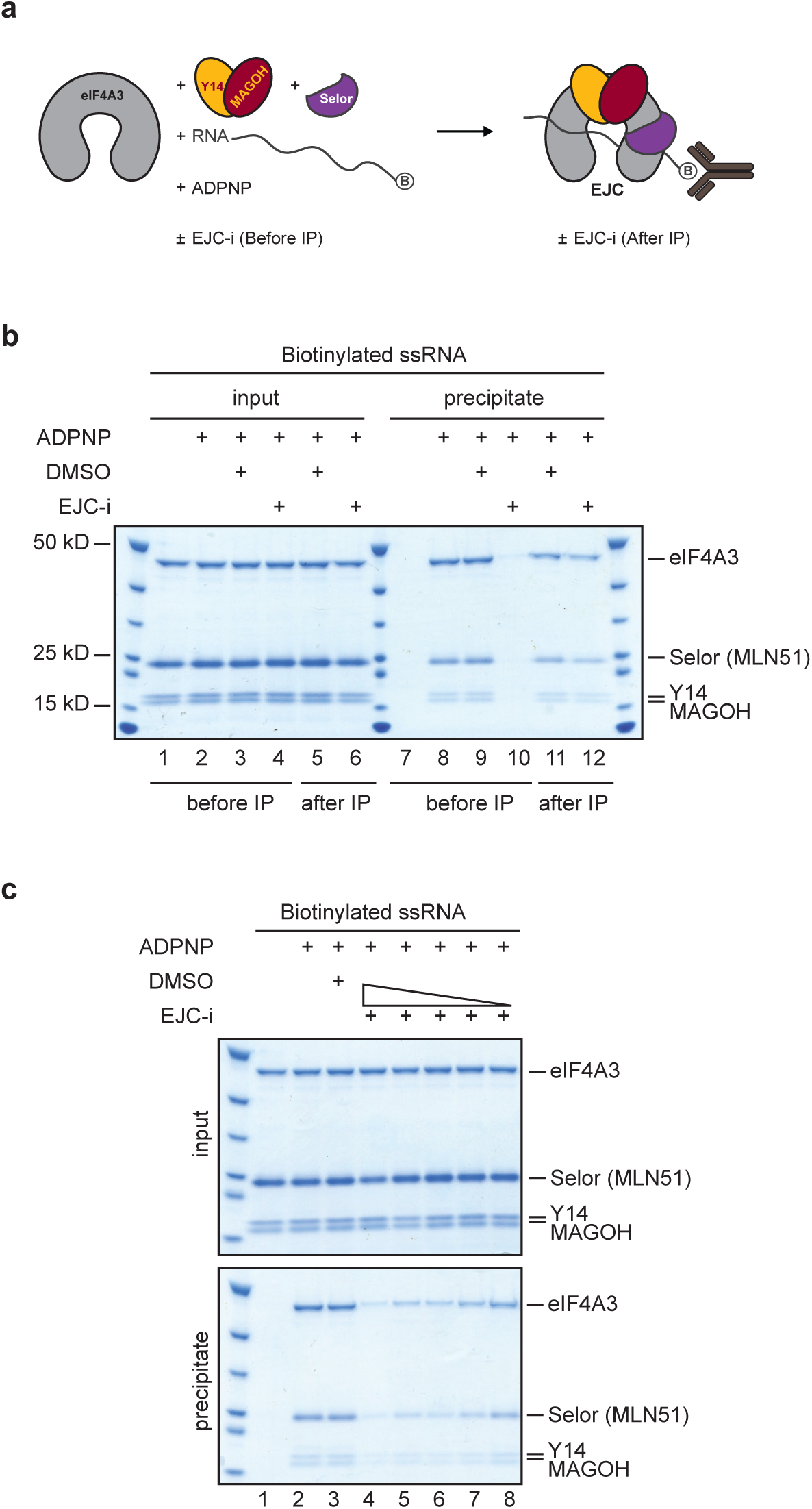
EJC-i inhibits *de novo* EJC assembly. **a** Scheme depicting the EJC reconstitution assay. **b** Protein coprecipitation with biotinylated ssRNA. Recombinant eIF4A3, MAGOH, Y14, and Selor were mixed with 3’-end biotinylated 30-mer ssRNA and incubated with or without ADPNP, in the presence or absence of 20 µM EJC-i or DMSO before or after coprecipitation, as indicated. Proteins from input (16% of total) and precipitate following denaturing elution were separated on a 4-12% (w/v) acrylamide SDS-PAGE in parallel with a protein marker. Protein molecular weight (kD) are indicated on the left, identity of proteins on the right. Gel was stained with Coomassie blue. **c** Same as (**b**) but incubation before coprecipitation was with decreasing doses of EJC-i (20 µM lane 4, 10 µM lane 5, 5 µM lane 6, 2 µM lane 7, 1 µM lane 8).

These reconstitution experiments clearly revealed that EJC-i blocks EJC assembly, but not *in vitro* pre-assembled EJCs. In order to confirm this observation, we tested the effect of the inhibitor on cell lysates that contain EJCs assembled in live cells (14). Cellular lysates from EJC split-luciferase expressing cells were incubated for 2 hours with different doses of EJC-i or DMSO as control. Even at the highest concentration of 20 µM, we did not observe any effect of EJC-i on luciferase activity (Supplementary Figure 2d). These results together strongly indicate that EJC-i efficiently inhibits *de novo* EJC assembly without affecting the stability of pre-assembled complexes.

### EJC-i affects alternative splicing and nonsense-mediated mRNA decay (NMD)

Recently, we edited cell lines with CRISPR/Cas9 to fuse a dTAG degron (39) to the EJC core subunits Y14 (24) and eIF4A3 (this work). An efficient EJC subunit depletion was obtained within a 24-hour time course (24) (Supplementary Figure 3). However, both the siRNA and degron knockdown approaches target a single protein and do not necessarily reflect the behavior of the entire complex, as they lead to degradation of both assembled and non-assembled EJC proteins. Furthermore, several hours to days of treatment are required. To assess the extent of inhibition of EJC functions *in vivo*, we monitored two established EJC-dependent functions, splicing regulation and NMD, looking at known target genes regulated by the activity of the complex, and compared HEK293T cells treated with a 20 µM of EJC-i for 24 hours to HEK293T cells carrying either the Y14 or the eIF4A3 dTAG degron depleted for 16 hours.

The EJC globally impacts splicing, notably by contributing to the recognition of neighboring introns and by suppressing cryptic splice sites usage (16, 20, 40–42). Through interaction with peripheral factors, including the ASAP and PSAP complexes, the EJC modulates different splicing choices and depletion of its core components or of any partners results most often in the exclusion of cassette exons. We analyzed transcripts originating from the *MRPL3*, *SMARCB1*, *GLRX3*, *SDHA*, and *HERC4* genes which generate mis-spliced forms when individual EJC core proteins are down-regulated (16, 20, 41). For all five genes tested, loss of EJC activity resulted in exon skipping events leading to accumulation of mis-spliced forms, both upon Y14 or eIF4A3 dTAG down regulation or upon EJC-i treatment (Figure 3a-e, compare lanes 1-3-5 to 2-4-6).

**Figure 3.**
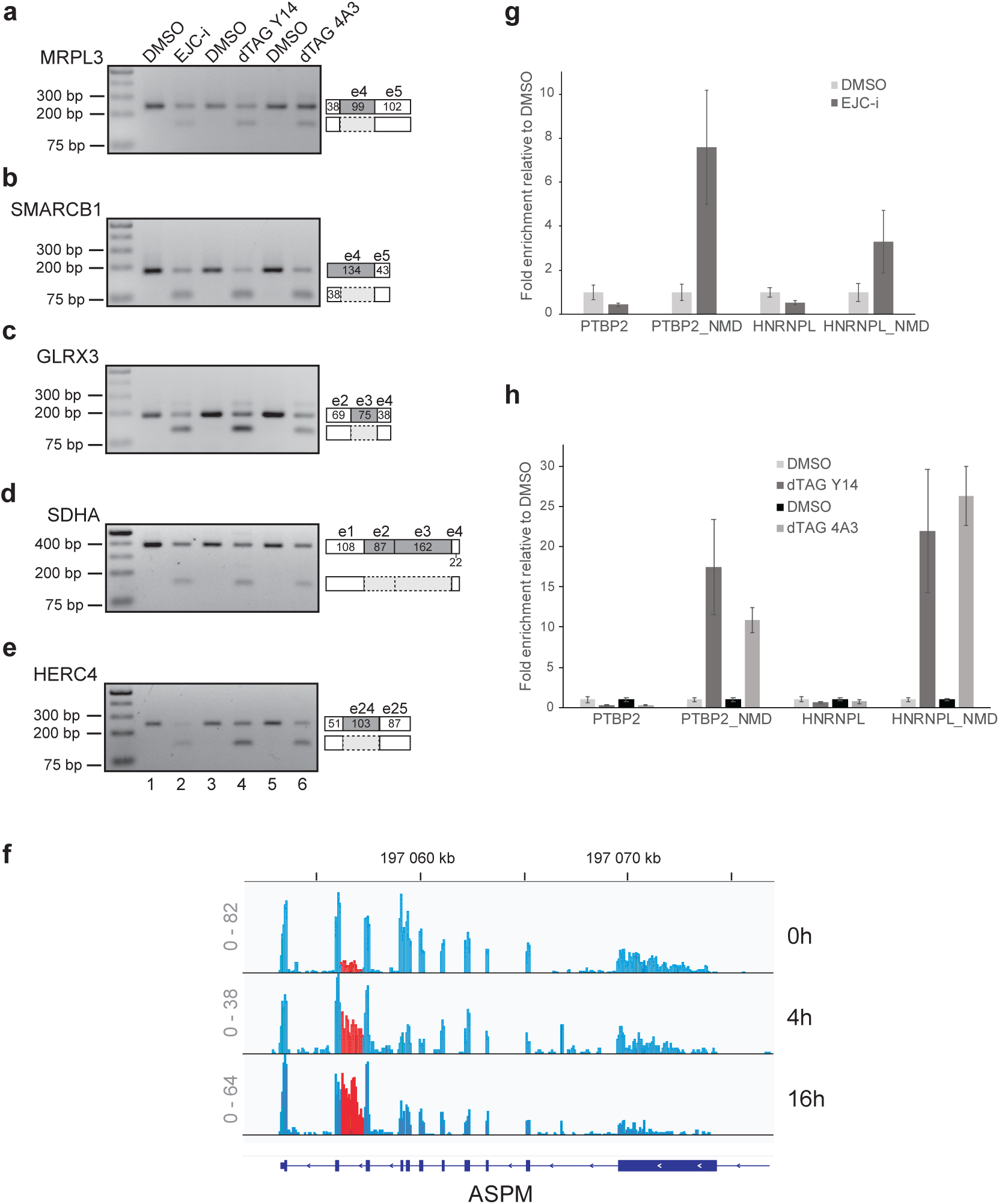
EJC-i affects alternative splicing and nonsense-mediated mRNA decay (NMD) **a** Endpoint RT-PCR analysis of *MRPL3* exon 4 skipping with RNA from HEK293T wild-type cells (lanes 1-2) or stably expressing a Y14 (lanes 3-4) or eIF4A3 (lanes 5-6) dTAG degron treated with either DMSO, 20 µM EJC-i for 24 hours, or dTAG for 16 hours, as indicated. cDNA was analyzed on 2% agarose gels stained with EtBr along with a DNA ladder. Sizes (bp) are indicated on the left. *MRPL3* exon architecture and alternatively spliced isoforms are schematized on the right. **b** Same as (**a**) for *SMARCB1* exon 4 mis-splicing. **c** Same as (**a**) for *GLRX3* exon 3 skipping. **d** Same as (**a**) for *SDHA* exon 3 and 4 skipping. **e** Same as (**a**) for *HERC4* exon 24 skipping. **f** IGV profile for *ASPM* mRNA at 0, 4, and 16 hours of 20 µM EJC-i treatment. Read scales are shown on the left. The retained intron is labelled in red. **g** RT-qPCR analysis of the *PTB2* and *HNRNPL* mRNAs and their NMD-sensitive isoforms from HEK293T wild-type cells treated with either DMSO or 20 µM EJC-i for 24 hours. **h** Same as (**g**) with RNA from HEK293T stably expressing a Y14 or eIF4A3 dTAG degron treated with either DMSO or dTAG for 16 hours.

Notably, inspection of RNA-seq data of EJC-i-treated cells over a 16-hour time-course (see below) revealed that the *ASPM* mRNA also strongly accumulates a mis-spliced form retaining the first-to-last intron (Figure 3f and Supplementary Figure 4a). A similar behavior was observed upon dTAG depletion of Y14 (Supplementary Figure 4b).

We next analyzed the effect of EJC-i on the control of NMD, for which the complex was originally identified (43–45). To evaluate the impact on EJC-dependent NMD activity, we performed RT-qPCR analysis on two specific genes, *PTBP2* and *HNRNPL*, that generate through splicing both stable mRNAs and unstable isoforms containing premature termination codons, and are therefore targeted by degradation through NMD (46). Following either EJC-i treatment (Figure 3g) or dTAG depletion (Figure 3h), the ensuing loss of EJC activity resulted in clear stabilization of NMD-sensitive isoforms. These results together show that chemical inhibition of EJC assembly and degron-mediated depletion of single complex core components have comparable effects on EJC-dependent regulation of alternative splicing and NMD.

### EJC-i prevents EJC-dependent exclusion of m6A deposition around splice junctions

We and others have recently unveiled a global function of the EJC in preventing m6A RNA modification around splice junctions (23–26). Using HEK293T cells carrying the Y14 dTAG degron, we have previously shown that Y14 depletion leads to transcriptome-wide increase of m6A within these EJC-mediated m6A exclusion zones (24). The physical presence of the complex may occlude methylation sites either by steric hindrance or generating an unfavorable mRNP architecture.

To demonstrate that these effects are indeed due to assembled EJCs, we performed an EJC-i treatment over a 16-hour time-course followed by transcriptome-wide m6A detection and quantification via m6A-seq2 (47), similar to our previous analysis with the Y14 dTAG degron (24). First, we calculated the m6A gene levels (m6A-GI). To capture the main trends of m6A gene level differences between the samples in an unbiased way, we performed a principal component analysis (PCA) for unbiasedly analyzing variability between samples and allowing for clear visualization and interpretation of underlying m6A level patterns. Principal component 1 (PC1) exhibited a clear increase over the samples, thereby capturing the increase m6A methylation level over the samples (Figure 4a). As in the Y14 dTAG degron m6Aseq2 experiment, we could observe that genes with multiple exons are affected in an exon-density-dependent way (24). This indicates that the m6A gene level for genes with more exons is stronger due to the lack of functional EJC-mediated m6A suppression (Figure 4b). In RNAs from EJC-i treated cells, a clear increase in m6A predominates in the exclusion zones 100-200 nucleotides around the exon-exon junction for long internal and last exons (Figure 4c). Furthermore, we performed *de novo* m6A peak calling based on all EJC-i-treated samples and found 31, 000 high confidence m6A sites (see Material and Methods). We calculated m6A-site scores across all samples and found that sites located in the EJC exclusion zones showed a significant increase in m6A methylation enrichment over the time course compared to other sites (Figure 4d). Overall, EJC-i treatment leads to a transcriptome-wide increase in m6A methylation and recapitulates observations obtained with the Y14 dTAG degron (24). So EJC-i rapidly inhibits the EJC in its m6A suppression properties, and reinforce the notion that impairing complex assembly specifically and effectively affects EJC-dependent m6A events on a global scale.

**Figure 4.**
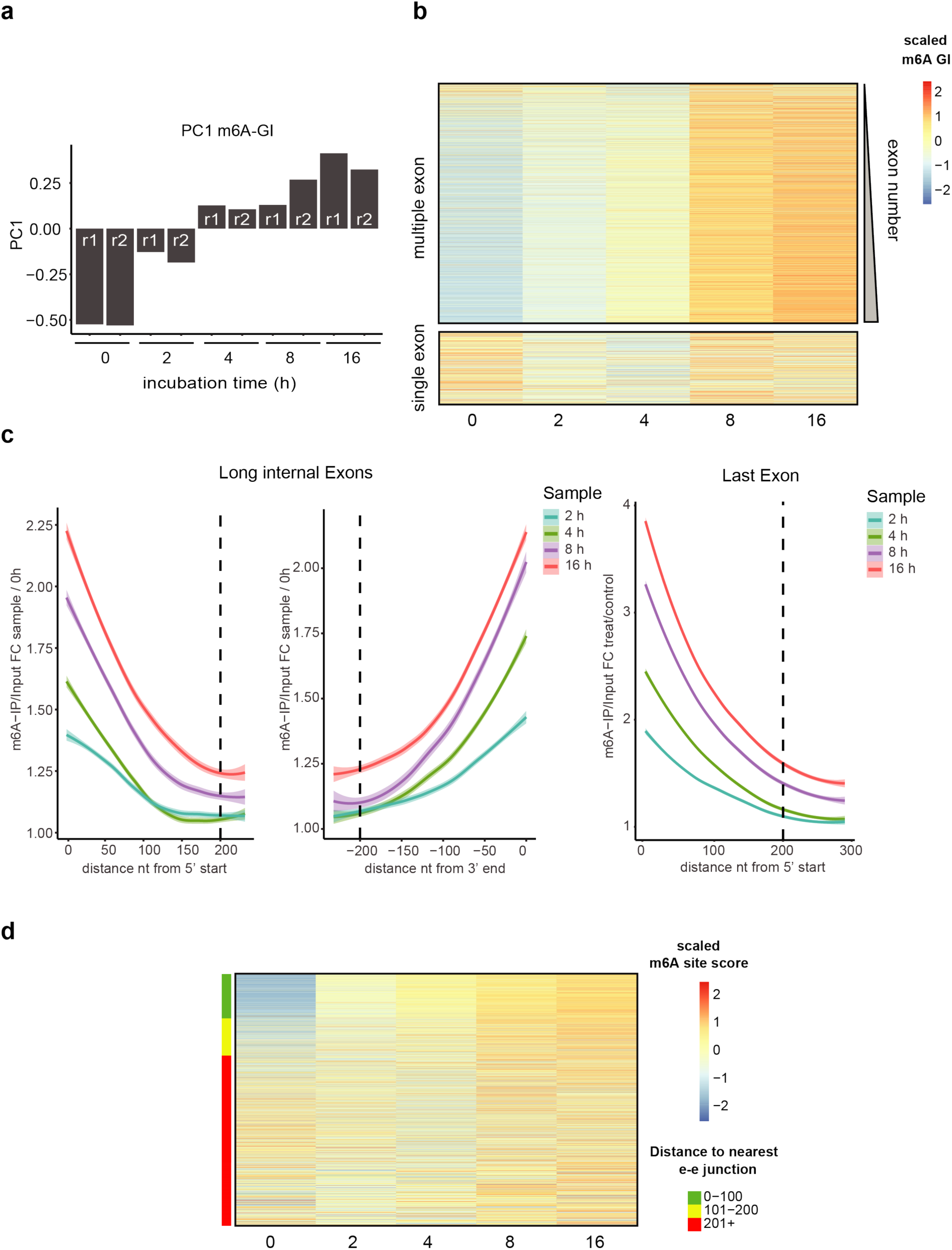
EJC-i prevents EJC-dependent exclusion of m6A deposition around splice junctions. **a** Barplot showing PC1 of principal component analysis of the measured m6A gene level (m6A-GIs) based on an m6A-seq2 experiment of an EJC-i time course experiment in HEK293T cells on two independent replicates (r1 and r2) **b** Heatmap showing the row-scaled m6A gene index (m6A-GIs) of all the samples of the EJC-i inhibition dataset for multiple exon genes (annotated with > one annotated exon, genes are ordered from lowest number of exons to highest, top) and single-exon genes (annotated with only a single annotated exon, bottom). **c** Meta fold-change of m6A enrichment (m6A-IP/Input) of timepoints after treatment of HEK293T cells with 20 µM EJC-i (2h, 4 h, 8 h, and 16 h) compared to timepoint 0h. Analysis was performed for 300 bases of the 5’ beginning of long internal exons (> 600 nt, left), for the last 240 bases towards the 3’ end (center), and for 300 nt from the beginning into the last exon (right). Calculated fold changes are based on the median m6A-IP/input score for 20-nt-long bins per time point. **d** Heatmap of row-scaled m6A site scores. High confidence m6A-sites are based on *de novo* peak calling (see Methods) of all EJC-i treatment and control samples. m6A-sites were ordered by the distance to the closest exon-exon junction.

### EJC-i promotes a mitotic arrest at prometaphase

EJC-i had been shown to arrest cells in G2M and to promote apoptosis (36). HEK293T cells stably expressing the Y14 dTAG degron were either depleted of Y14 through treatment with dTAG or assembly of the EJC blocked by treatment with 10 µM EJC-i over a 24-hour time course. Cell cycle profiles were assessed by flow cytometry after treatment and compared to DMSO control. Prolonged inhibition of EJC function by either method resulted in a clear cell cycle arrest at the G2/M boundary to a comparable extent, with 37.3% cells in G2/M following dTAG depletion of Y14 and 26.7% after EJC-i block (Figure 5a). Microscopic inspection of cells revealed that the arrest due to interfering with EJC stability corresponded to cells arrested in prometaphase (Figure 5b).

**Figure 5.**
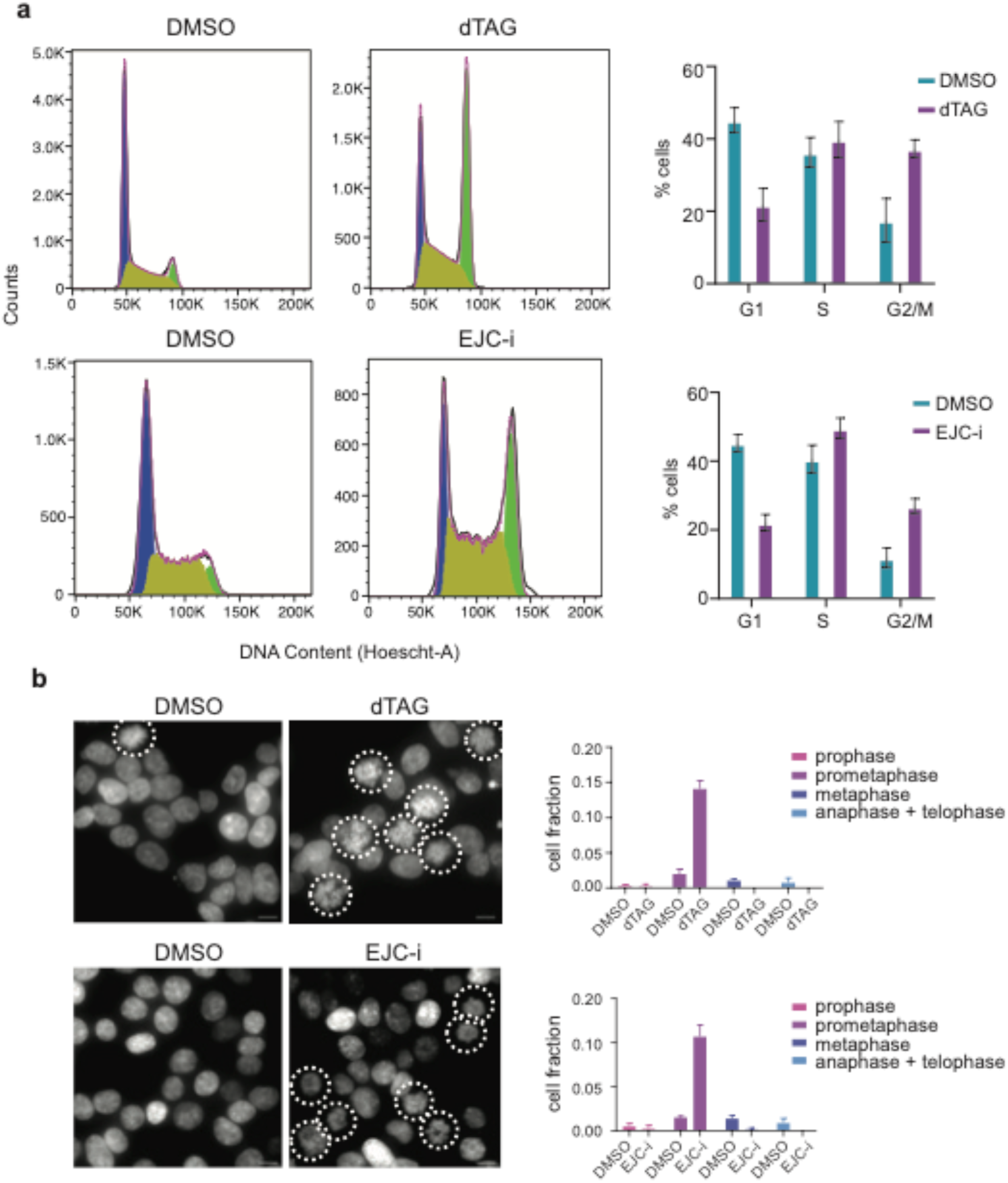
EJC-i promotes a mitotic arrest at prometaphase. **a** Cell cycle profiles (counts on vertical axes vs. DNA content on horizontal axes) were assessed by flow cytometry after treatment of HEK293T cells stably expressing the Y14 dTAG degron with DMSO, dTAG, or 10 μM EJC-i for 24 hours. The percentage of cells assigned to the respective stages of the cell cycle is indicated on the respective charts on the right. **b** Representative DAPI staining of the same cells as in (**a**) with fraction of cells assigned to the respective stages of the cell cycle quantified on the respective charts on the right. Dotted lines encircle cells arrested in prometaphase.

These results show that targeted knockdown of Y14 core EJC subunit and EJC-i similarly result in mitotic arrest of cell cycle at prometaphase.

### EJC-i prevents proper intracellular localization of mRNAs

In human cells, *NIN* mRNAs localization to centrosomes depends both on translation and the EJC, and siRNA-mediated EJC downregulation impairs pericentriolar material organization (22). Numerous transcripts are specifically transported to centrosomes at different stages of the cell cycle (22, 48–51). To explore in more detail the role of the EJC in their localization, we tested the effect of EJC-i. First, we looked at *NIN* mRNAs, using a HeLa cell line stably expressing a GFP tagged version of Centrin1 to label centrosomes (Figure 6). As previously reported, *NIN* mRNA decorated centrosomes during interphase and mitosis, but this localization was dramatically lost upon inhibition of EJC assembly by EJC-i (Figure 6a). We previously identified *ASPM* and *NUMA1* mRNAs among a set of 8 mRNAs localized at the centrosomes in HeLa cells, with *ASPM* mRNA localizing to centrosome specifically during mitosis (48–51). Both mRNAs displayed a clear localization to the centrosomes, which was abolished after treatment with EJC-i (Figure 6b, 6c, 6d). In addition, the same effect was observed for *PCNT* mRNAs, which also localize to centrosomes (48) (Supplementary Figure 5a), and for *AKAP9* mRNAs, which accumulate on the Golgi apparatus close to centrosomes (50) (Supplementary Figure 5b). These results identify a set of five transcripts whose localization in proximity to centrosome depends on assembly of the EJC, generalizing the effect previously reported for *NIN* mRNA. As the localization of these mRNAs also require ongoing translation, it points toward a localization mechanism in which the EJC and the nascent peptides work together to localize polysomes (22, 48, 51). Together with the observation that EJC knockdowns affect centrosome organization (22), these data strengthen the notion that the EJC complex is intricately involved in a post-transcriptional regulation program defining centrosomal functions during the cell cycle.

**Figure 6.**
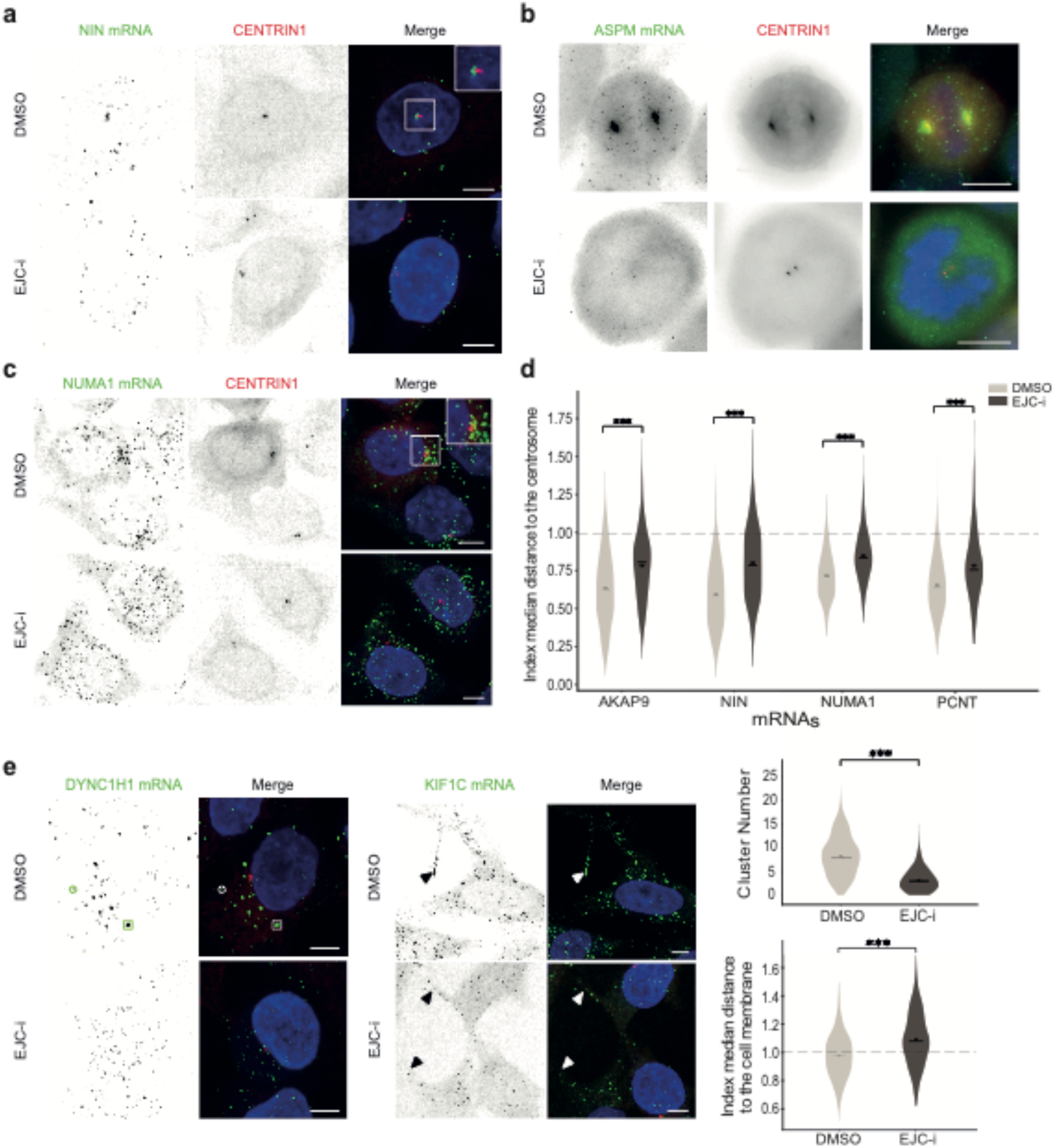
EJC-i prevents proper intracellular localization of mRNAs. **a** Images are micrographs of HeLa cells treated or not with EJC-i for 16h, and labelled for *NIN* mRNA by smFISH (left), and centrin1-GFP protein (middle). Merged image (right) display *NIN* mRNAs in green, centrin1 in red, and DNA in blue. Scale bars: 10 microns. **b** Same as (**a**) for *ASPM* mRNA. **c** Same as (**a**) for *NUMA1* mRNA. **d** Index of normalized median distances of mRNAs to centrosomes, for the two conditions, measured on individual cells. In each cell, distances from spots to closest centrosome were normalized using distances from a random RNA distribution, a distance inferior to one indicates colocalization whereas a distance superior to one indicates anti-colocalization. The dots represent the mean for all cells and the bar the median. P-values were determined using an Alexander Govern and a Games-Howell tests. *** P<0, 001. **e** Same as (**a**) for *DYNC1H1* mRNA (left) and *KIF1C* mRNA (middle). Right: plot of the quantification of the number of mRNA foci per cell, in cells treated or not with EJC-i for 16h (up) and plot of the median of the normalized distance of mRNAs to the cell membrane for cells treated or not with EJC-i for 16h (down). Circles indicate single molecule of mRNA, squares a foci and arrows indicate protrusions.

We next tested the impact of EJC-i on transcripts that localize to other cellular compartments (50). *DYNC1H1* code for the large subunit of dynein and its mRNAs accumulate into discrete cytoplasmic foci acting as translation factories (52), and *KIF1C* codes for a plus-end kinesin motor whose mRNAs concentrate into cytoplasmic protrusions (50). Remarkably, in both cases, exposure to EJC-i severely prevented their localization (Figure 6e). These results identify two additional transcripts whose localization is dependent on the function and integrity of the EJC.

The localization of the centrosomal mRNAs cited above is translation dependent, as is the formation of the *DYNC1H1* translation factories. We thus tested the effects of EJC-i on the translation of these transcripts. We previously generated HeLa cells having a SunTag fused at the N-terminus of the endogenous *ASPM* and DYNC1H1 proteins, effectively labeling their polysomes and nascent peptides (48, 52). After treatment with EJC-i, we observed a drastic reduction in the number of *ASPM* polysomes in interphase cells, already visible after 4h of treatment and complete after 16h (Figure 7a). Interestingly, the export of *ASPM* mRNAs was also inhibited. Its mRNAs molecules remained in the nucleus after EJC-i addition, accounting for the absence of polysomes in the cytoplasm (Figure 7a). This effect of the EJC-i on nuclear export was specific for *ASPM* mRNAs as it was not seen for *DYNC1H1* transcripts, which were exported and translated similarly whether EJC-i was present or not (Figure 7b). Next, we checked *ASPM* translation in mitotic cells. The SunTag-ASPM protein localized at the centrosome and spindle whether EJC-i was present or not (Figure 7c). As noted above, the *ASPM* mRNA did not localize to centrosomes in EJC-i treated cells. Interestingly however, some active SunTag-ASPM polysomes were observed in EJC-i treated cells, suggesting that the SunTag-*ASPM* mRNA could be translated after its release from the nucleus, following the breakdown of the nuclear envelope. Altogether, these data indicate that EJC-i can inhibit the export of specific mRNAs and that its effects on mRNA localization were not due to an inhibition of translation. These data point to the advantageous use of EJC-i as a tool to efficiently monitor the involvement of the EJC in mRNA localization.

**Figure 7.**
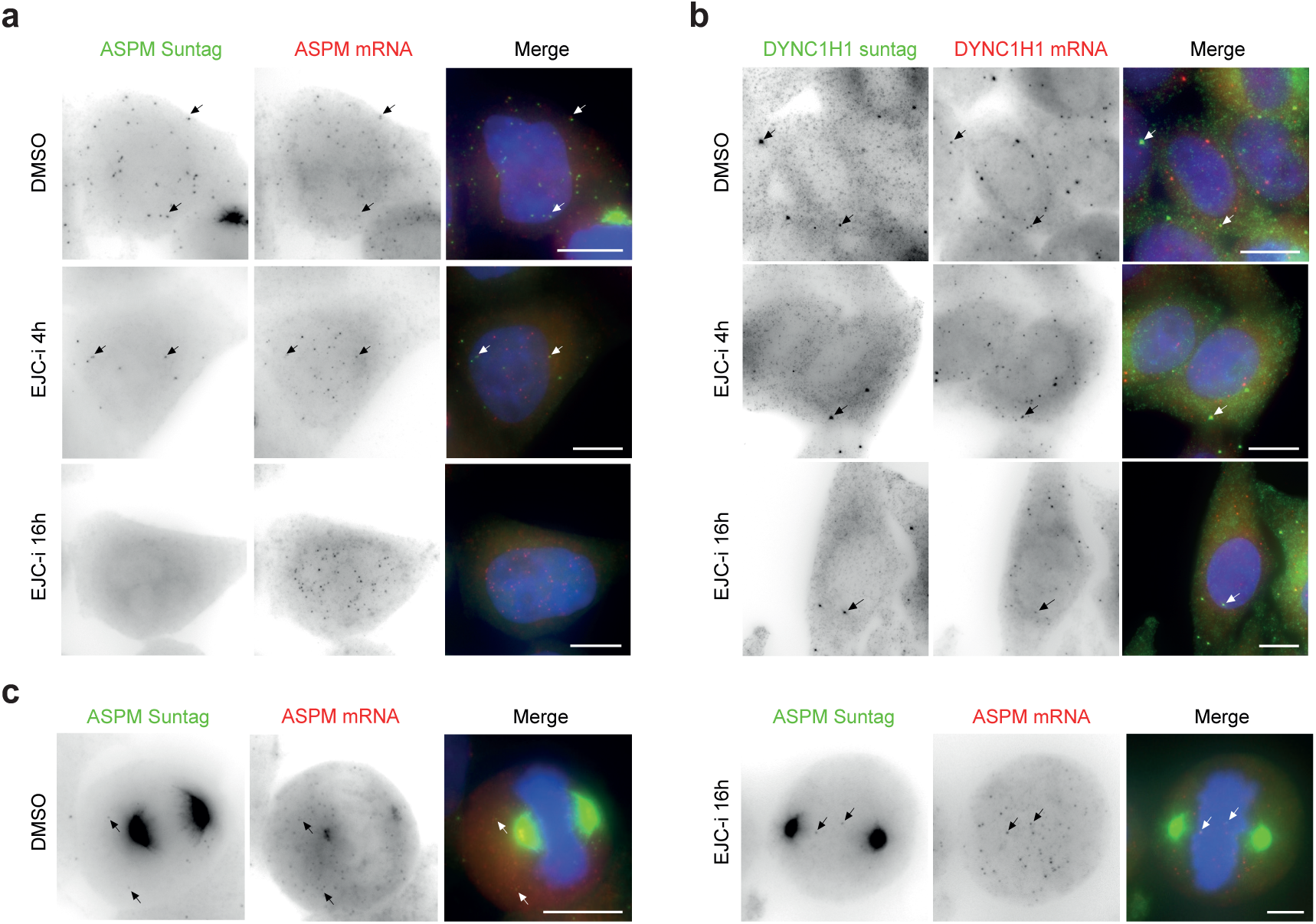
Selective inhibition by EJC-i reveals the EJC-dependent export of ASPM mRNAs. **a** Images are micrographs of HeLa cells, in interphase, treated or not with EJC-I for 4h or 16h, expressing a SunTag-ASPM allele (left) SunTag-ASPM protein display in green, ASPM mRNAs in red and DNA stained with DAPI in blue. Right: same as left with cells expressing a SunTag-DYNC1H1. Scale bars: 10 microns. Arrows indicate polysomes. **b** Same as (**a**) for SunTag-*ASPM* proteins and mRNA for cells in mitosis.

### EJC-i disrupts ciliogenesis in mouse Neural Stem Cells

The integrity of centrosome organization is necessary for the differentiation of radial glial mouse Neural Stem Cells (NSC). When these quiescent mono-ciliated cells are starved for serum, they become quiescent and differentiate into multi-ciliated ependymal cells that populate the surface of brain ventricles, ensuring the flow of cerebrospinal fluid through the beating of their cilia (53). In NSC, the primary cilium forms from the centrosome’s mother centriole (basal body), while during differentiation, amplification of centrioles leads to the generation of multiple motile cilia in ependymal cells (54). We previously reported that EJCs accumulate at basal bodies of quiescent NSC and RPE1 cells (22). siRNA downregulation of EJC core components results in abnormal ciliogenesis and decrease in the number of ciliated RPE1 cells (22). However, this procedure is challenging with NSC primary cultures.

This prompted us to analyze the effect of EJC inhibition on ciliogenesis in primary cultures of NSC isolated from newborn mouse forebrain (54). Cultures were treated with 10 µM EJC-i or control DMSO at the onset of ependymal cell differentiation and over two days. Treatment with the inhibitor led to a marked reduction in the primary cilium length as well as to delayed differentiation (Figure 8a, 8b). Thus, ciliogenesis is severely disrupted upon inhibition of EJC in NSC cells like in RPE1 cells, confirming its crucial role in this central process.

**Figure 8.**
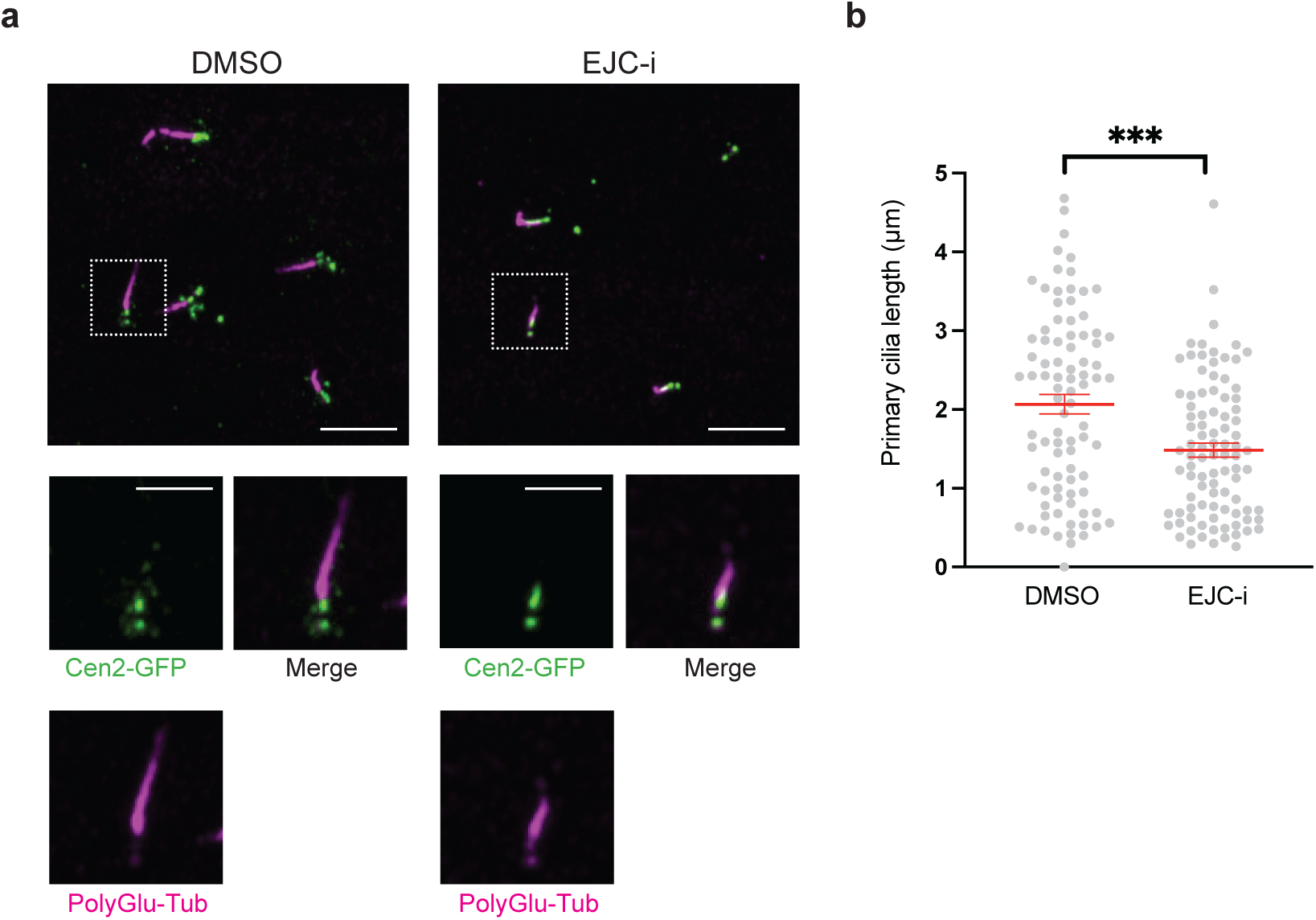
EJC-i disrupts ciliogenesis in mouse Neural Stem Cells. **a** Representative images of NSC treated with 10 µM EJC-i or DMSO as control. Primary cilia were stained by poly-glutamylated tubulin (PolyGlu-Tub) antibody (magenta), centrioles by Centrin2-GFP transgenic mice (green). Scale bars in the upper and lower panels are 5 and 2, 5 µm, respectively. **b** Quantification of the difference in primary cilia length between the two conditions. The bars represent the mean +/- SEM; one dot corresponds to one cell (3 biological replicates). P values were determined using a two-sided Student’s t-test. *** P < 0, 001.

## DISCUSSION

Here, we provide evidence that a compound known to inhibit eIF4A3 ATPase activity inhibits *de novo* EJC assembly, without affecting pre-formed complexes. We show that this drug impairs known EJC-dependent functions such as alternative splicing, NMD, m6A deposition, and mitotic progression. Its effects are similar to degron-mediated downregulation of the core EJC components eIF4A3 and Y14. Use of the drug supports the EJC-dependent localization of several centrosomal mRNAs. Inhibition of EJC assembly disrupts ciliogenesis in mouse NSC primary cultures, further substantiating a central role of the EJC in centrosome biology. Moreover, we propose a broader function of the EJC in the transcript targeting to other cell compartments.

EJC-i targets a key component of the EJC complex without affecting the expression level of its essential subunits. It had been described as a chemical compound that selectively inhibits the ATPase activity of eIF4A3 (33, 35, 36). However, ATPase activity is not required for EJC assembly (38, 55). Binding of EJC-i appears to occur within an allosteric region of eIF4A3 wherein some of the most responsive stretches either include or are in close proximity with residues involved in interactions with RNA and MAGOH (9, 10, 55). Noteworthy, we find that EJC-i impairs interaction of eIF4A3 with RNA in the absence of its partners. EJC-i binding might induce a conformational change in eIF4A3 that prevents these interactions and thereby EJC assembly (33). Together with a prevalence of cytosolic material extracted by standard methods for total protein preparation, the impact of EJC-i on eIF4A3 co-precipitation with the other core factors is hard to observe by Western blot. However, the effect of EJC-i *in vivo* can be followed quantitatively using an EJC-NanoBiT assay (14). Consistent with recent observations that EJCs slowly disassemble in vivo, it plateaus at about 80% inhibition, likely reflecting the presence of complexes not undergoing, or only slowly, recycling. Exposure to EJC-i leads to the same alterations in alternative splicing, m6A deposition, centrosome structure, mitotic delay and ciliogenesis defects than targeted degradation of Y14, another EJC subunit. Thus, although eIF4A3 may have a life independently of the EJC (56, 57) and other EJC-independent effects cannot be excluded, EJC-i can be considered primarily as an EJC inhibitor. Blocking EJC complex assembly provides a unique opportunity to pinpoint events depending upon functional EJCs.

Many mRNAs display a specific intracellular localization. The EJC was initially found to play a key role in the localization of *oskar* mRNA in Drosophila oocytes (21). Here we show that several other human mRNAs are affected by EJC-i. *NIN* mRNA had already been reported to require the EJC to be localized at centrosomes in mammalian cells (22). We now extend this requirement to other centrosome-localized transcripts such as *ASPM*, *NUMA1* and *PCNT*, as well as *AKAP9* mRNAs which accumulate on the Golgi apparatus close to centrosomes. Furthermore, localization of *DYNC1H1* mRNA in translation factories and of *KIF1C* mRNA in cytoplasmic protrusions also require the EJC. Interestingly, among these mRNAs affected by EJC-i, localization can occur in either translation-dependent or independent ways, indicating the involvement of the EJC in multiple RNA transport mechanisms. While localization of *NIN*, *NUMA1*, *ASPM, AKAP9* and *DYNC1H1* requires translation, that of *PCNT* and *KIF1C* mRNAs does not (22, 48, 50, 51). In the case of *DYNC1H1* mRNAs, EJC-i did not induce a shutdown of their translation, pointing toward a direct effect of the EJC in mRNA localization. In the case of *ASPM* mRNAs, EJC-i blocked export from the nucleus at interphase, resulting in nuclear accumulation and a lack of cytoplasmic mRNAs. *ASPM* mRNA export might have a particularly strong requirement in the human Transcription Export (TREX) complex that associates with the EJC (19, 58), and/or be the consequence of the observed intron retention upon EJC assembly inhibition (Figure 3f and Supplementary Figure 4). However, these nuclear *ASPM* mRNAs are released into the cytoplasm at mitosis when the nuclear envelope breaks down, they become translated, yet they fail to localize to centrosomes in EJC-i treated cells. This observation points towards a coordinated role for the EJC and local translation in mRNA localization. The EJC and the nascent peptides might work together to localize the corresponding polysomes at the right place. Noteworthy, the EJC is associated with stalled ribosomes (14). Arrested ribosomes having already synthesized N-terminal fragments of proteins, might direct transport of the corresponding polysomes to their destination (48).

EJC-i leads to a mitotic arrest at prometaphase. This is likely linked to the impaired centrosome organization due to EJC inactivation. Indeed, the centrosome and its above-mentioned components play a key role in mitosis. Noteworthy, knockdown of Ninein leads to aberrant spindle formation (59). ASPM regulates mitotic spindle orientation during metaphase (60). NUMA protein is an essential player in mitotic spindle assembly and maintenance (61). AKAP9 is required for association of the centrosomes with the poles of the bipolar mitotic spindle during metaphase (62). Therefore, impaired localization of mRNAs coding for these proteins might be responsible for the mitotic arrest.

Both EJC core and peripheral components have been implicated in various cancers (27). Furthermore, hypomorphic expression of eIF4A3 and Y14 has been linked to two neurodevelopmental genetic disorders in humans: Richieri-Costa Pereira syndrome and Thrombocytopenia with absent radius syndrome, respectively (28, 63, 64). Mouse models have established a clear connection between imbalances in EJC core components and defects in neural stem cell (mNSC) division, neurogenesis, and cortical development (65, 66). We had previously shown that Y14 or eIF4A3 siRNA knockdown in a retinitis pigmentosum epithelium model cell line (RPE1) impairs centrosome organization and ciliogenesis (22). A centrosome forms the basal body of cilia (67). The defective ciliogenesis might be due to the defective local synthesis of critical centrosome components. The use of siRNA with mNSC is challenging. Thanks to EJC-i, we now show that EJC is also essential for ciliogenesis in primary mNSCs. It might be hypothesized that EJC-dependent mitotic defects are responsible for mNSC division.

Popular methods for the inhibition of essential proteins rely on conditional gene knockouts using CRISPR/Cas9 genome editing or conditional knockdown methods such as shRNA- or siRNA or by dTAG degron modules introduced by genome editing (39). While effective, conditional loss of function of essential genes have several limitations. Typically, RNA silencing methods necessitate extended time (days) to achieve successful downregulation and suffer from potential off-target and side effects. Not all cell types are amenable to CRISPR/Cas9-based knockdown or knockout approaches. This is indeed the case for mouse NSC primary cultures. Small molecules that specifically target essential proteins to inhibit or modulate their function circumvent these limitations. Their action is rapid, generally limited by diffusion into cells, they are easily applicable in diverse settings without the necessity for genetic alterations, and carry the potential for therapeutical applications (68, 69). In conclusion, EJC-i is the first available specific inhibitor of the EJC as a whole. It opens up new venues to investigate the roles that the complex plays in the regulation of cell cycle and consequences of its dysfunction associated with pathologies and developmental defects.

## DATA AVAILABILITY

The high throughput RNA sequencing data are available in the GEO database under accession number: GSE286119

## SUPPLEMENTARY DATA

Supplementary Data are available at NAR online.

## AUTHOR CONTRIBUTIONS

O.B., H.L.H., and T.V. conceived the project. T.V. performed NanoBiT experiments, in vitro reconstitution assays, immunoprecipitations and Western blotting, endpoint PCRs and RT-qPCRs analyses. D.D. and S.S. performed the m6A sequencing and their computational analyses. C.B. performed cell cycle analysis and established the eIF4A3 degron cell line. L.G. and C.B. performed flow cytometry experiments. O.P., F.S., and E.B. performed immunofluorescence microscopy, smFISH, SunTag experiments, and image analysis including quantifications. M.F. and N.S. prepared and conducted analysis on primary cell cultures of mNSC. T.V. wrote the paper that was reviewed and edited by O.B. and H.L.H.

## ACKNOWLEDGEMENTS

We thank all HLH lab members for fruitful discussions and for critical reading of the manuscript.

