## Supplemental figures for "Chemical inhibition of Exon Junction Complex assembly impairs mRNA localization and neural stem cells ciliogenesis"

**Supplementary Figure 1**

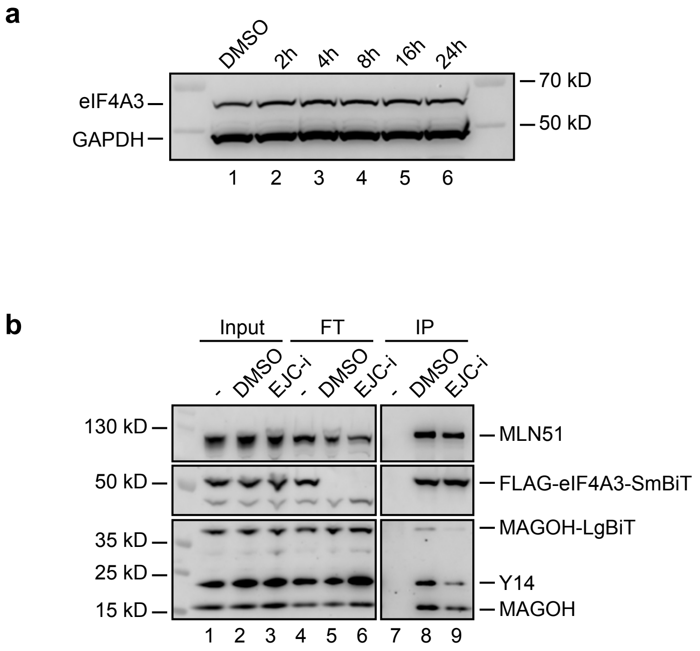


**Supplementary Figure 1. (related to Figure 1) EJC-i decreases the amounts of EJC in live cells**

**a** Western blot analysis of the indicated proteins from HEK293T cells treated with DMSO or a 20 µM dose of EJC-i for the indicated times (hours). **b** Western blot analysis of EJC proteins coimmunoprecipitated (IP) with anti-FLAG from protein extracts of HEK293T cells co-expressing MAGOH-LgBiT and FLAG-eIF4A3-SmBiT treated with either DMSO or a 20 µM dose of EJC-i for 4 hours. Lanes labelled - represent no antibody controls. FT is the flow-through fraction.

**Supplementary Figure 2**

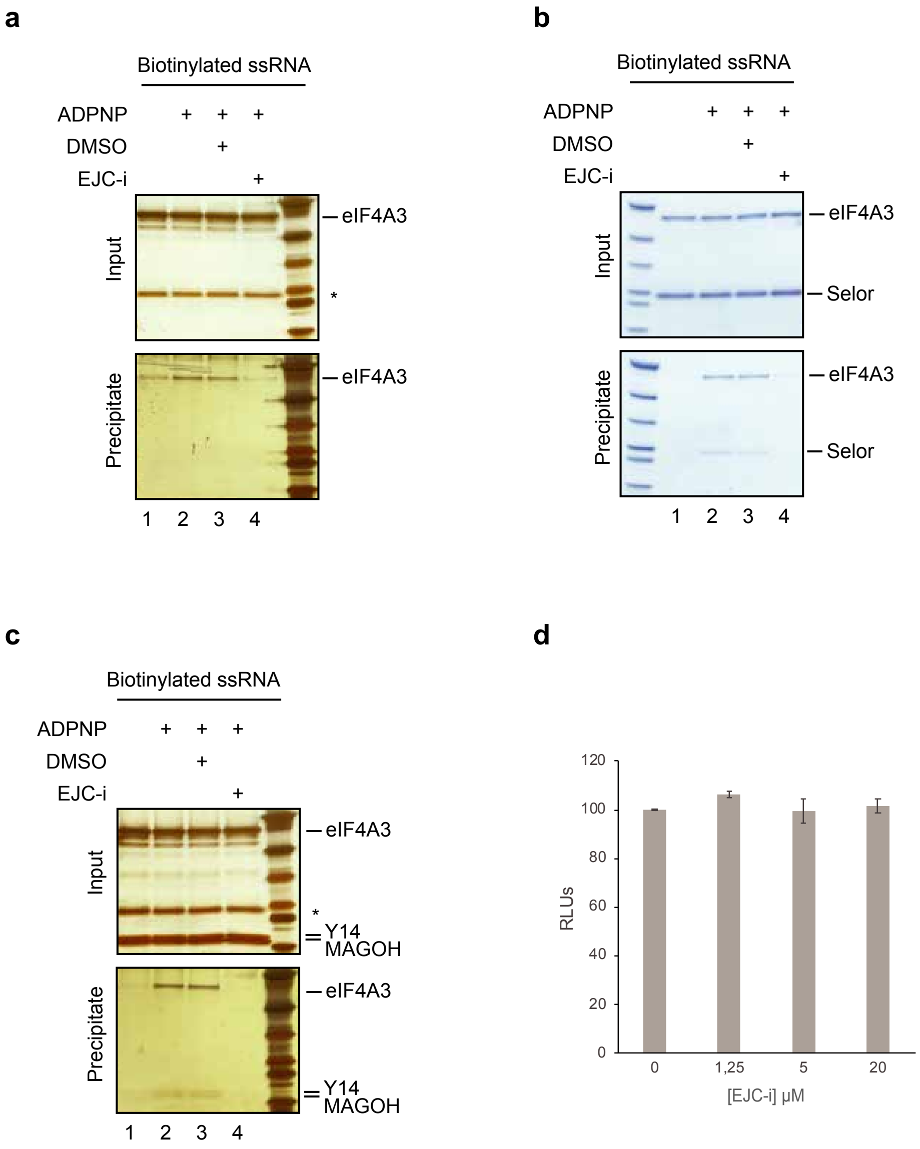


**Supplementary Figure 2. (related to Figure 2) EJC-i inhibits *de novo* EJC assembly**

**a** Coprecipitation with biotinylated ssRNA. Recombinant eIF4A3 was mixed with 3'-end biotinylated 30-mer ssRNA and incubated with or without ADPNP, in the presence or absence of 20 µM EJC-i or DMSO before coprecipitation. Proteins from Input (16% of total) and Precipitate following denaturing elution were separated on a 4-12% (w/v) acrylamide SDS-PAGE in parallel with a protein marker. Gel was stained with silver. Position of proteins is indicated on the right. Asterisk marks an eIF4A3 breakdown product. **b** Same as (**a**) with eIF4A3 and Selor. Gel was stained with Coomassie blue. **c** Same as (**a**) with eIF4A3 and the MAGOH/Y14 heterodimer. Gel was stained with silver. Asterisk marks an eIF4A3 breakdown product. **d** Response of a lysate from a stable HEK293T cell line co-expressing MAGOH-LgBiT and FLAG-eIF4A3-SmBiT to a 2 hours incubation with the indicated µM doses of EJC-i expressed as relative luciferase units (RLUs).

**Supplementary Figure 3**

a

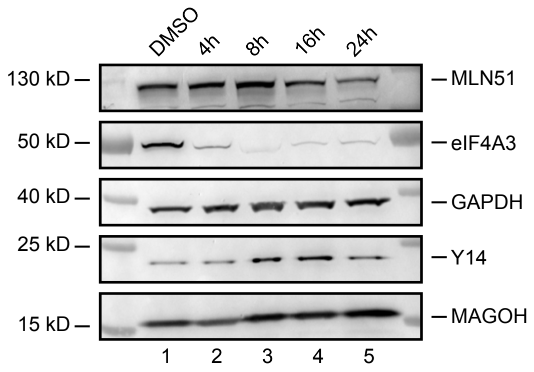


b

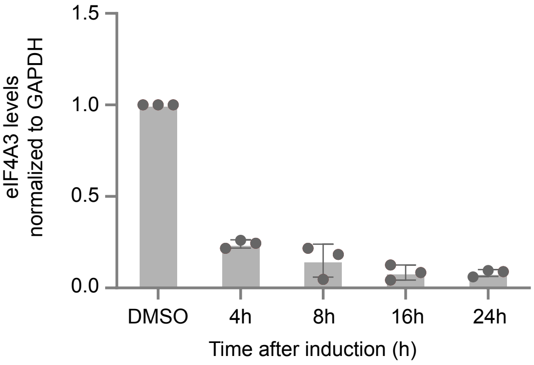


**Supplementary Figure 3. (related to Figure 3) EJC-i affects alternative splicing and nonsense-mediated mRNA decay (NMD)**

**a** Representative Western blot images of eIF4A3 degron and control DMSO-treated time points, with antibodies against different EJC members and control GAPDH. **b** Quantification of Western blot band intensities of eIF4A3 normalized to GAPDH in the eIF4A3 degron time points, based on three biological replicates. Grey dots mark individual measurement data points, error bars indicate standard deviation of the mean.

**Supplementary Figure 4**

a

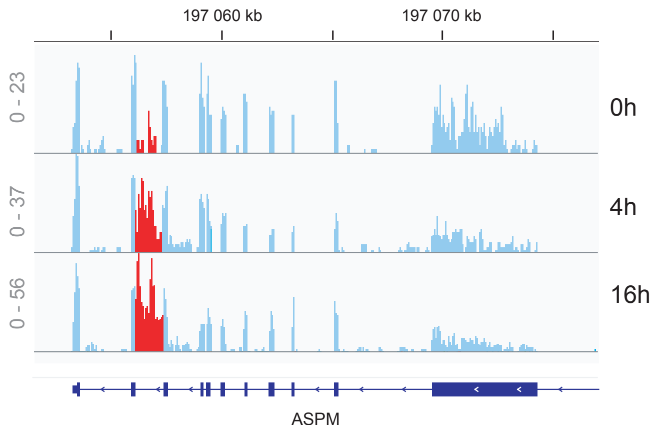


b

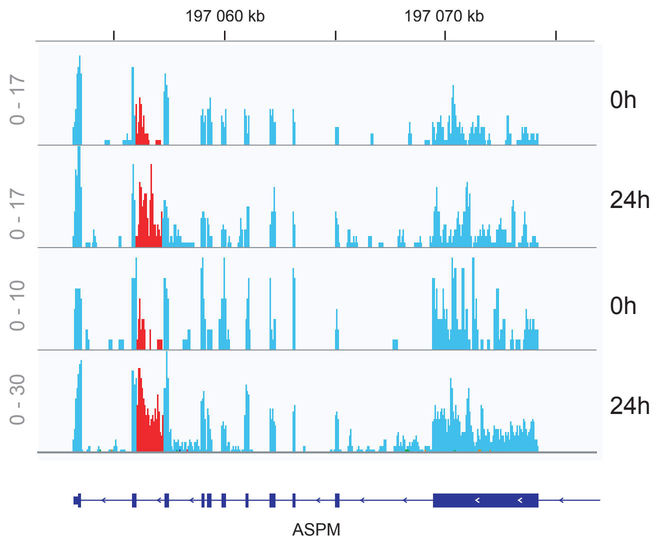


**Supplementary Figure 4. (related to Figure 3) EJC-i affects alternative splicing and nonsense-mediated mRNA decay (NMD)**

**a** IGV profile for a second biological replicate of *ASPM* mRNA at 0, 4, and 16 hours of 20 µM EJC-i treatment. Read scales are shown on the left. The retained intron is labelled in red. **b** IGV profile for two biological replicates of *ASPM* mRNA at 0 and 24 hours of dTAG-mediated Y14 knockdown. Read scales are shown on the left. The retained intron is labelled in red.

**Supplementary Figure 5**

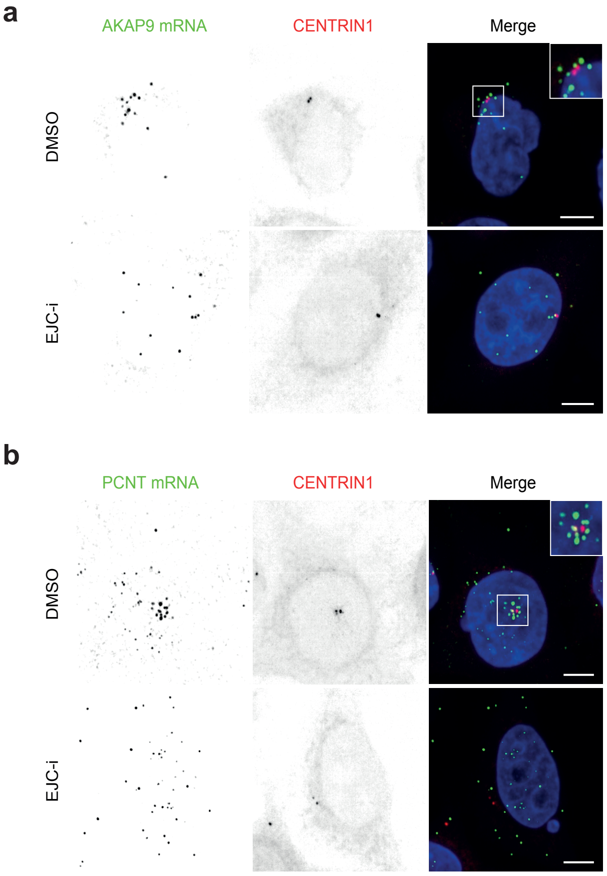


**Supplementary Figure 5. (related to Figure 6) EJC-i prevents proper intracellular localization of mRNAs**

**a** HeLa cells stably expressing Centrin1-GFP after smFISH treated with 10 µM EJC-i for 16 hours or DMSO as control. Fluorescent signal corresponding to *AKAP9* mRNA (left), to the Centrin1-GFP protein (middle), and merged image (right) with mRNA signal in green, Centrin1 in red, and DNA stained with DAPI in blue. Scale bars: 10 microns. **b** Same as (**a**) for *PCNT* mRNA.
